# Mitochondrial Dynamics Modulation via MyoAAV-Mediated OPA1 Overexpression or BNIP3 Knockdown in Pancreatic Ductal Adenocarcinoma (PDAC)-Induced Cachexia

**DOI:** 10.64898/2026.11.13.766453

**Authors:** Ruqaiza Muhyudin, Nicole N. Noga, Francielly Morena, Sydney Hilgenbrink, Muskan Taori, Allie Ivy, Maryam Khaledabadi, Mary S. Jia, Christopher Nelson, Tyrone Washington, Nicholas P. Greene

**Affiliations:** Cachexia Research Laboratory, Exercise Science Research Center, Department of Health, Human Performance and Recreation, University of Arkansas, Fayetteville, Arkansas, United States; Cell and Molecular Biology Graduate Program, University of Arkansas, Fayetteville, Arkansas, Fayetteville, Arkansas, United States; Department of Physiology and Aging, University of Florida, Gainesville, Florida, United States; Department of Biomedical Engineering, University of Arkansas, Fayetteville, Arkansas, Fayetteville, Arkansas, United States; Exercise Muscle Biology Laboratory, Exercise Science Research Center, Department of Health, Human Performance and Recreation, University of Arkansas, Fayetteville, Arkansas, United States

**Keywords:** Cancer Cachexia, skeletal muscle, mitochondrial quality control, fusion, mitophagy

## Abstract

Disruption of mitochondrial dynamics and quality control precedes overt muscle wasting in cancer cachexia, but whether modulating it specifically in muscle in the presence of systemic wasting could preserve tissue mass and function remain elusive. We tested two independent, but complementary approaches in the orthotopic preclinical model of pancreatic ductal adenocarcinoma (KPC); inducing inner-mitochondrial membrane fusion through OPA1 overexpression and suppressing mitophagy via BNIP3 knockdown. Male and female C57BL/6J mice received bilateral intramuscular tibialis anterior (TA) injections of myotropic adeno-associated virus (MyoAAV) encoding murine either OPA1 cDNA or a BNIP3-targeting short hairpin RNA two weeks prior to orthotopic implantation of KPC^1245^ pancreatic cancer cells. Raw body and tissue weights, *in vivo* dorsiflexor torque and fatigability, TA fiber size and type, protein fractional synthesis rate, oxidative histochemistry, and selected molecular markers were assessed at the humane endpoint. Neither intervention prevented systemic wasting, however, in males, OPA1 overexpression partially attenuated TA mass loss, relieved torque deficits, and restored the proportion of type IIB fibers. Females developed a milder cachectic phenotype; OPA1 overexpression increased TA mass and shifted fiber composition toward type IIA fibers without any changes in torque. In males, BNIP3 knockdown markedly improved torque, restored fatigue resistance, preserved type IIB fiber proportion, and lowered DRP1 protein. Functional benefits concurred by BNIP3 shRNA delivery were accompanied by lower mixed-protein synthesis in both sexes. These findings support mitochondrial quality control as a therapeutic target while indicating that effective treatment will require calibrated, rather than maximal, suppression of mitochondrial turnover.

**NEW & NOTEWORTHY:** The effects of muscle-directed modulation of mitochondrial quality control using MyoAAV on contractile function in pancreatic cancer cachexia remain unclear. We show that MyoAAV mediated OPA1 overexpression or BNIP3 knockdown yield distinct contractile benefits along with higher type IIB fibers proportion. These findings identify muscle mitochondria as potential target using translationally relevant approach to mitigate cachexia-associated weakness and highlight a functional benefit.

## INTRODUCTION

Cancer cachexia (CC) is a systemic and complex metabolic syndrome characterized by progressive and involuntary loss of skeletal muscle mass, with or without fat loss that cannot be reversed through nutritional supplementation alone (1). This syndrome affects up to 80% of cancer patients with advanced stage malignancies and is responsible for approximately 20-30% of cancer related deaths (2, 3). Pancreatic ductal adenocarcinoma (PDAC) carries one of the highest cachexia burdens among all cancer types, with muscle loss directly contributing to increased surgical complications, reduced treatment tolerance, diminished physical function and compromised quality of life (4–6). Despite significant efforts we currently lack FDA-approved effective therapies to prevent and reverse cachexia and thus further work is needed to identify best therapeutic targets. A growing body of evidence indicates mitochondrial dysfunction as an early and rate-limiting mechanism in cachexia related skeletal muscle atrophy (7), preceding measurable muscle mass loss (8). Specifically, excessive fission and impaired fusion capacity coupled with excessive mitophagy, are primary processes contributing to the onset and progression of muscle atrophy in different muscle pathologies (9) including cancer cachexia (10–14). Despite increasing evidence linking mitochondrial dysfunction and degeneration with CC, it remains unclear whether targeting mitochondrial dynamics directly in skeletal muscle can preserve mass and function in PDAC cachexia and whether such a response is sex specific.

OPA1 (Optic Atrophy 1) is a dynamin related GTPase responsible for the fusion of inner mitochondrial membrane and cristae remodeling and is reduced progressively in atrophic muscle (15, 16). Prior studies reported OPA1 overexpression to result in significant protection against denervation induced muscle loss through stabilized cristae architecture of mitochondria along with suppressed apoptotic signaling (17). Most importantly, our laboratory recently demonstrated that whole-body transgenic OPA1 overexpression attenuates tumor induced muscle loss and preserves contractile force in Lewis lung carcinoma (LLC) tumor bearing male and female mice (18). Conversely, BNIP3 (BCL2/adenovirus interacting protein 3), a BH3-only protein functioning as a selective autophagy cargo receptor (19), is reported to be progressively upregulated in cachectic muscle across multiple preclinical models (20–22). Our recent findings show that global BNIP3 deletion failed to protect muscle mass in cancer induced cachexia in LLC mice despite improving mitochondria quality (23). In turn, BNIP3-null animals demonstrated suppressed basal protein synthesis under non-tumor control conditions (23), further linking mitochondrial status to muscle proteostasis. Therefore, it is likely global deletion of BNIP3 preceding cancer-induced stress created an unfavorable energetic environment leading to a potentiated proteostatic deficit. These collective findings suggest spatial and temporal precision in modulating mitochondrial dynamics is critical for effective therapeutic intervention. Muscle specific and partial suppression of BNIP3 expression may contribute to preserve basal mitophagy dependent mitochondrial quality control which can further protect skeletal muscle mass with the reduction of pathologic excess driven by aberrant FoxO3 hyperactivation (24). Although transgenic models have been used extensively to provide mechanical insights, viral-vector mediated delivery represents a pragmatic advancement toward clinical applicability with more translational value.

Adeno-associated virus mediated gene transduction may represent an optimal opportunity for targeted therapy. Specifically, MyoAAV, a variant AAV capsid, has up to 80-fold greater transduction efficiency in skeletal muscle with less off target expression as compared to parental AAV9 (25) while the recent approval of AAV-based gene therapies by the FDA for neuromuscular disorders establishes clinical precedent for AAV-mediated gene therapy (25–30). MyoAAV, thus, presents a promising platform to investigate muscle intrinsic therapeutic targets in CC as being investigated earlier as well (31, 32).

The present study was designed to test the hypothesis that muscle directed modulation of mitochondrial dynamics through MyoAAV mediated delivery of murine OPA1 complimentary DNA or BNIP3 targeting short hairpin RNA, preserves skeletal muscle mass and function during PDAC-induced cachexia. This preventive intervention paradigm differs from previous studies with post-onset therapeutic approaches, as it allows mechanistic interrogation of whether mitochondrial quality control at the onset of a catabolic challenge determines the trajectory of muscle wasting. This study demonstrated the role of mitochondrial remodeling in preserving muscle function relative to muscle mass in sex specific manner.

## MATERIAL AND METHODS

### Animals and Housing

Male and female C57BL/6J mice (Jackson Laboratory; 000664, Bar Harbor, ME, USA) were used in all experiments and housed in temperature and humidity-controlled conditions with a 12-hour light/dark cycle. Mice were provided *ad libitum* access to standard laboratory chow (LabDiet: 5053 - PicoLab® Rodent Diet 20) and water for the duration of the study. All animal procedures were approved by the University of Arkansas Institutional Animal Care and Use Committee (IACUC) and adhered to the NIH guidelines for Care and Use of Laboratory Animals. Both biological sexes were included in experiments as a variable and mice were randomly divided in experimental groups (n=9-12 per group per sex, specifics detailed below). Implantation of pancreatic cells (KPC FC1245) was performed on animals at 12 weeks of age around the time of 8:00-11:30 a.m. (∼1-4 hours into their light cycle) and animals were monitored daily for body weight and behavior to indicate any stress or pain. Tissue collection was performed at pre-determined humane endpoint (n= 6) or experimental endpoint (∼19 days post tumor implantation for males and ∼17 days post implantation for females; n= 78). Humane endpoints were determined by either ∼20% body weight loss or moribund behavior. Tissue collection was performed between 8:00 and 13:00 h.

### MyoAAV Vector Construct Design and Production

For this study, two different MyoAAV constructs were designed. For OPA1 overexpression, a plasmid was constructed to encode murine OPA1 complementary DNA along with transcriptional control of human creatine kinase promoter (33). For reporter, the OPA1 cDNA was primarily tagged with 13xMyc tag at the carboxy terminus. For BNIP3 knockdown (experiment 2), a plasmid was engineered to encode distinct short hairpin RNA sequences (shRNA) to target non-overlapping regions of murine BNIP3 mRNA under U6 polymerase III promoter. The shRNA sequence was derived from Sigma-Aldrich MISSION pLKO-puro construct (clone NM_009760.2-524s1c1; TRCN0000009691), with the target sequence 5′-GCATGAGAAACACAAGCGTTA-3′ (34). The plasmid was formulated as a self-complementary vector to enhance expression (33). For reporter gene, blue fluorescent protein (BFP) was co-expressed driven by a CAG promoter. Whole plasmid sequencing was used to verify both plasmids’ constructs. Plasmid constructs and MyoAAV capsid were sent to University of North Carolina Vector Core for MyoAAV vectors production. Vectors were received with the yield of 1.8^13^ vg/ml (OPA1-AAV) and 2.6^11^vg/ml (BNIP3 shRNA-AAV).

### Intramuscular AAV Vector Administration

Mice received intramuscular injections of MyoAAV at 10 weeks of age directly in tibialis anterior muscles bilaterally, two weeks prior to tumor implantation. Mice were anesthetized via oxygen mixed isoflurane inhalation (5% induction and 2% maintenance). Briefly, the hair was shaved, and anterior leg was aseptically prepared. MyoAAV or saline (as sham control) (35, 36)was administered at standardized volume (40 µl) at distal and proximal parts of the TA muscle (2^10^vg/TA for OPA1-AAV, 2^9^vg/TA for BNIP3 shRNA-AAV).

### Orthotopic KPC Tumor Model

Two weeks following intramuscular MyoAAV administration, mice (12 weeks of age) underwent surgery for orthotopic KPC^1245^ pancreatic cell implantation. KPC cells (FC1245) were received from Cold Spring Harbor Laboratory (courtesy of Dr. David Tuveson). Cells were maintained in DMEM supplemented with 10% fetal bovine serum and 1% penicillin-streptomycin. Cells were trypsinized at 90% confluency, passage number: 2-3, washed twice in PBS and suspended in 50 µl (150,000 viable cells) media (Matrigel: PBS, 1:1). KPC cell suspension was injected directly to the pancreas using a 29-guage needle (37). Control groups received vehicle (Matrigel: PBS, 1:1). All animals were monitored daily for body weight (BW) loss (not to exceed 20% loss in body weight from the initial BW (day of KPC injections)) and behavioral indicators of stress. Buprenorphine was injected subcutaneously (0.1mg/kg), before tumor implantation Meloxicam tablets (Bio-Serv; MD275-0125) were given to animals for 72 hours for post-surgery pain management. Same group of animals included in control and KPC Sham (Cancer: KPC, AAV: Saline) groups were used in both interventions to compare them with their sex matched intervention group (KPC OPA1(OE), BNIP3(KD)).

### In-vivo Muscle Contractile Function

Animals were assessed for contractile function of their dorsiflexor muscles, 48h before tissue collection, as previously described (23). Briefly, Mice were anesthetized (oxygen mixed isoflurane at 5%) and left hindlimb was shaved and cleaned with ethanol pad for the procedure. Animals’ body temperature was maintained to 37°C using heating pad. The foot was placed and held in a footplate using surgical tape. On either side of the peroneal nerve, two electrodes (Model E2–12, Grass Technologies, RI, USA) were inserted percutaneously, and instant stimulation was used to ensure the correct placement of electrodes. After confirmation from instant stimulation, muscle contraction was induced via electrical stimulation through electrical stimulator and stimulation isolation unit (Model 300B-LR, Aurora Scientific, Aurora, Ontario, Canada). Torque was measured during 12 isometric contractions at various stimulation frequencies (10, 20, 30, 40, 60, 80, 100, 125, 150, 200, 250 and 300 Hz). The purpose of using a range is to determine different contractile phenotypes as physiological hindlimb motor recruitment happens at 60Hz, and 100Hz frequencies and low frequency fatigue phenotype can be determined at supramaximal stimulation frequencies such as >200 Hz (38, 39). The muscle was then subjected to fatigability test after 4-minute break in between by applying 120 isometric contractions at 40 Hz with the duration of 150ms for the total time duration of 2 minutes. Ambulatory behavior was closely monitored during the recovery time period.

### Protein Fractional Synthesis Rate (FSR)

Protein synthesis was determined using gas chromatography-mass spectrometry (GC-MS) (5977A and 7890A Agilent, Santa Clara, CA, US) by calculating fractional synthesis rate (FSR) performed as described (23, 40). Mice were given a bolus of 99.9% deuterium oxide (D₂O) (151882-1L, Millipore Sigma, Burlington, MA, USA) intraperitoneal injection at 20 µL/g of their body weight 24 hours prior to tissue collection, and drinking water was then supplemented with 4% D20 for the remaining time period, in order to maintain the plasma pool of D2O. Blood was collected via cardiac puncture at the time of tissue collection in tube with heparin to avoid coagulation and plasma was isolated via centrifugation (1500 x g, 20 minutes, 4°C) in order to calculate D2O plasma pool. Approximately 10 mg of TA tissue was powdered, homogenized and washed in 10% Trichloroacetic acid (TCA) solution in order to discard any cytosolic amino acids and proteins. 6 M HCl (300 µl) was added to the final pellet and heated at 100°C overnight to hydrolyze proteins into amino acids. 50 µL of hydrolysate was then transferred into another tube to dry it completely in order to further derivatize amino acid pool with solution of methyl-8, methanol and acetonitrile (3:2:1, v/v). The final solution was then transferred to GC-MS capillary column (Agilent; 5182-0549) and placed in the GC-MS sample rack. GC-MS ran sample at the volume of 1 µL with 20:1 split ratio. Plasma was used to determine precursor pool of ^2^H_2_O and was treated with 5% acetone solution in acetonitrile and 10M NaOH overnight at room temperature in order to conjugate the available ^2^H_2_O to acetone. The acetone was extracted by using n-Hexane solution and finally placed in GC-MS capillary column to be analyzed at 20:1 split ratio. A ratio of deuterated alanine over undeuterated alanine was used to access protein synthesis in tissues, while ratio of conjugated acetone over acetone for precursor pool determination. Mixed protein’s FSR was calculated by the equation (EA * [EBW * 3.7 * t(h)] – 1 * 100) (10, 11), where EA is the amount of protein bound deuterated alanine, EBW represents the quantity of precursor pool of ^2^H_2_O in the body and t(h) is the time in hours the label was present in the body (40).

### Immunoblotting

Immunoblotting was performed as described by our laboratory (18). Frozen TA muscle samples were powdered and approximately 15 mg of tissue was suspended in 200 µl of 2X protein sample buffer [50 mM Tris-HCl, 1% sodium dodecyl sulfate (SDS), 10% glycerol, dithiothreitol (20 mM), 2-mercaptoethanol (127 mM), and 0.01% bromophenol blue combined with protease inhibitors, and phosphatase inhibitors at pH 6.8] and homogenized. Samples were then boiled for 5 minutes at 95°C, centrifuged for 5 minutes at maximum speed and supernatant was collected for protein quantification via RC/DC assay (5000122, BioRad, Hercules, CA). 40ug of total protein was loaded in 10% gel and resolved by SDS-PAGE at 165V for 50 minutes, transferred to PVDF membrane at 25V for 35 minutes using semi-dry transfer system (Bio-Rad Trans-Blot Turbo Transfer System (1704150EDU). Ponceau stain was applied to membrane after transfer step to visualize and scan the protein bands. The membranes were then incubated with blocking solution (LICOR bio; 927-60001) for 1 hour at room temperature on rocking machine. The membranes were washed with Tris-buffered saline (TBS) 3×5 minutes and then followed by overnight incubation with primary antibody solution (1:1000 dilution) in LICOR blocking buffer and TBS with 0.2% Tween 20. All primary antibodies were from Cell Signaling Technology. The membranes were incubated at 4°C to antibodies optic atrophy 1 (OPA1) (80471), cytochrome c oxidase subunit 4 (COXIV) (4844), dynamin-related protein 1 (DRP1) (14647S) and BCL2 interacting protein 3 (BNIP3) (3769). Following overnight incubation, membranes were washed with TBS 3 times for 5 minutes and further incubated with respective secondary antibodies (IRDye 800CW Goat anti-Rabbit IgG Secondary Antibody, IRDye 800CW Goat anti-Mouse IgG Secondary Antibody) for 1 hour at room temperature. They were then washed with TBST 3 times for 5 minutes, followed by image scan on Li-Cor Odyssey FC via IR detection. Target protein band intensities were normalized to 45-KDa Ponceau S-stained band as a loading control (41). Samples were normalized to an internal control loaded on all gels to account for gel-to-gel variability.

### RNA isolation, cDNA synthesis and Quantitative Real-time PCR

RNA isolation, cDNA synthesis and quantitative real-time PCR were performed as previously described by our laboratory (10, 11, 42). Frozen TA muscle samples (∼10 mg) were powdered and suspended in 1 ml of TRIzol reagent (Fisher Scientific; 15596018) and homogenized using Bead Mill 24 homogenizer (Fisherbrand, Thermo Fisher Scientific, Waltham, MA) with following settings: 2 cycles of 15s each, displacement (D)= 0.10 and speed (S) =6. 200 µL of chloroform was added to the homogenized solution followed by 3 minutes incubation at room temperature for phase separation and then 25 minutes centrifugation at 4°C. After centrifugation, the clear top portion of the solution was carefully transferred to another tube followed by addition of equal volume of 70% of diethyl pyro carbonate (DEPC)-treated ethanol for RNA precipitation. The solution was then added to RNAeasy column, with further processing of sample through RNA isolation kit (ThermoFisher Scientific; Catalog #12183025). RNA was collected in 20 µL of RNAase free water and RNA concentration and purity was determined by fluorometry using 260/280nm ratios (> 1.8) read on Bio-Tek Power Wave XC microplate reader (BioTek Instruments Inc., Winooski, VT) with Take3 microvolume plate using Gen5 software. VILO superscript reagent was used to reverse transcribed RNA into cDNA and then diluted to 1:100 (10ng/ µl). Quantitative real-time PCR was performed using TaqMan probe reagent and QuantStudio 3 PCR instrument (Applied Biosystems, Waltham, MA) with the following setting: Denaturation at 95°C for 20 seconds, 40 cycles of annealing and extension of 95°C for 1 second and 60°C for 20 seconds. The final quantification of mRNA level was calculated using the ΔΔCT method, from the cycle threshold (C_t_) values. Target TaqMan probes for *18s* (Mm03928990_g1), *Deptor* (also referred as *depdc6;* Mm01195339_m1), *FBXO32* (Mm00499523_m1), *Trim63* (Mm01185221_m1), *Ddit4* (hereafter referred to as *Redd1*; Mm00512504_g1), *Ncam1* (Mm01149710_m1), *Runx1* (Mm01213404_m1), *Ryr1* (Mm01175211_m1), *Atp2a1* (Serca1) (Mm01275320_m1) and *Atp2a2* (Serca2; Mm01201431_m1) were purchased from Thermo Fisher Scientific Gene Expression assays. All targets were normalized to the 18s C_t_ value, which did not differ between experimental groups. Genes are italicized and proteins are not.

### Immunofluorescence staining

Another set of muscle cross-sections from sham, OPA1 (OE) and BNIP(KD) groups were used for reporter gene immunofluorescence microscopy (43, 44). Sections were air dried, lined with hydrophobic pen barrier and then incubated overnight at 4°C with primary antibodies for dystrophin (1:100, ab15277, Abcam, St. Louis, MO, USA) and anti-Myc tag antibody (1:100, ab32, Abcam, St. Louis, MO, USA). For BNIP3 (KD) experiment, sections were fixed with 4% paraformaldehyde for 10 minutes after sectioning followed by incubation with primary antibody for dystrophin (1:100, ab15277, Abcam, St. Louis, MO, USA) overnight at 4°C. Following incubation, slides were washed with PBS for three times and then incubated with isotype-specific secondary antibody cocktail of 1:200 donkey-anti-rabbit IgG (Alexa Fluor 647, magenta) and 1:200 goat-anti-mouse IgG1 (Alexa Fluor 488, green; only for OPA1 (OE) group) in PBS at room temperature for 75 minutes in dark. Following by three rounds of PBS washes, slides were then mounted using ProLong Diamond Antifade Mountant (ThermoFisher Scientific; P36961).

### Succinate Dehydrogenase Histological Staining (SDH), Cross-sectional area (CSA) and Fiber Typing

SDH staining and immunohistochemistry was performed as described by members our laboratory previously (8, 43). One of the TA muscles from each animal was snap frozen in liquid nitrogen-cooled isopentane for all cryosectioning and staining procedures. Each section was cut at 8 μm onto positively charged slides using a Leica CM1860 cryostat (Leica Biosystems, Wetzlar, Germany). Sections were air dried for 1 hour, lined with hydrophobic pen barrier and then first stained for succinate dehydrogenase (SDH). Sections were incubated in incubation media (50 mM sodium succinate, 50 mM phosphate buffer, 0.12 M KH_2_PO_4_, and 0.88 M Na_2_PHO_4_, 0.5 mg/mL nitroblue tetrazolium) for 60 minutes at room temperature. Following incubation, slides were washed with dH_2_O 3 times and then proceeded for Fiber type staining. Sections were incubated with primary antibodies for dystrophin (1:100, ab15277, Abcam, St. Louis, MO, USA), MyHC IIA (1:100 SC-71, Developmental Studies Hybridoma Bank, Iowa City, IA, USA), and MyHC IIB (1:100 BF-F3, Developmental Studies Hybridoma Bank, Iowa 151 City, IA, USA) were applied overnight in a PBS cocktail at 4°C. Following incubation, slides were washed with PBS for three times and then incubated with isotype-specific secondary antibody cocktail of 1:200 donkey-anti-rabbit IgG (Alexa Fluor 647, magenta), 1:200 goat-anti-mouse IgG1 (Alexa Fluor 488, green), and 1:200 goat-anti-mouse IgM (Alexa Fluor 555, red) in PBS at room temperature for 75 minutes protected from light. Following three rounds of PBS washes, slides were then mounted using ProLong Diamond Antifade Mountant (ThermoFisher Scientific; P36961).

### Imaging and Analysis

Zeiss Axiolmager M2 was used to image whole sections with Zen software at 20x magnification (Carl Zeiss Microscopy, Oberochen, Germany). Fibers intensity for SDH assay, Fiber cross-sectional area and fiber type was quantified using the semiautomated software MyoVision (45, 46). In order to ensure accurate quantification, post-analysis manual corrections of the regions that were damaged or folded were performed for their removal. The analyst was blinded to experimental groups during these assessments.

### Statistical Analysis

All statistical analysis was performed using SAS 9.4 (SAS institute Inc. NC, USA). The OPA1 over expression and BNIP3 knockdown experiments were analyzed as independent studies and males and females were analyzed separately, although control and KPC Sham group animals were shared between both experiments to compare them to respective interventions. Results with only two relevant groups (Fig 1C, 1D, 1F and 1G) were analyzed using an unpaired two-tailed student t-test. Other outcomes (involving three groups) were analyzed using one-way analysis of variance (ANOVA). When the overall ANOVA was significant, pairwise comparisons were performed for between group significance using Tukey-Kramer adjustment for multiple comparisons. Statistical significance was set at p < 0.05.

**Figure 1:**
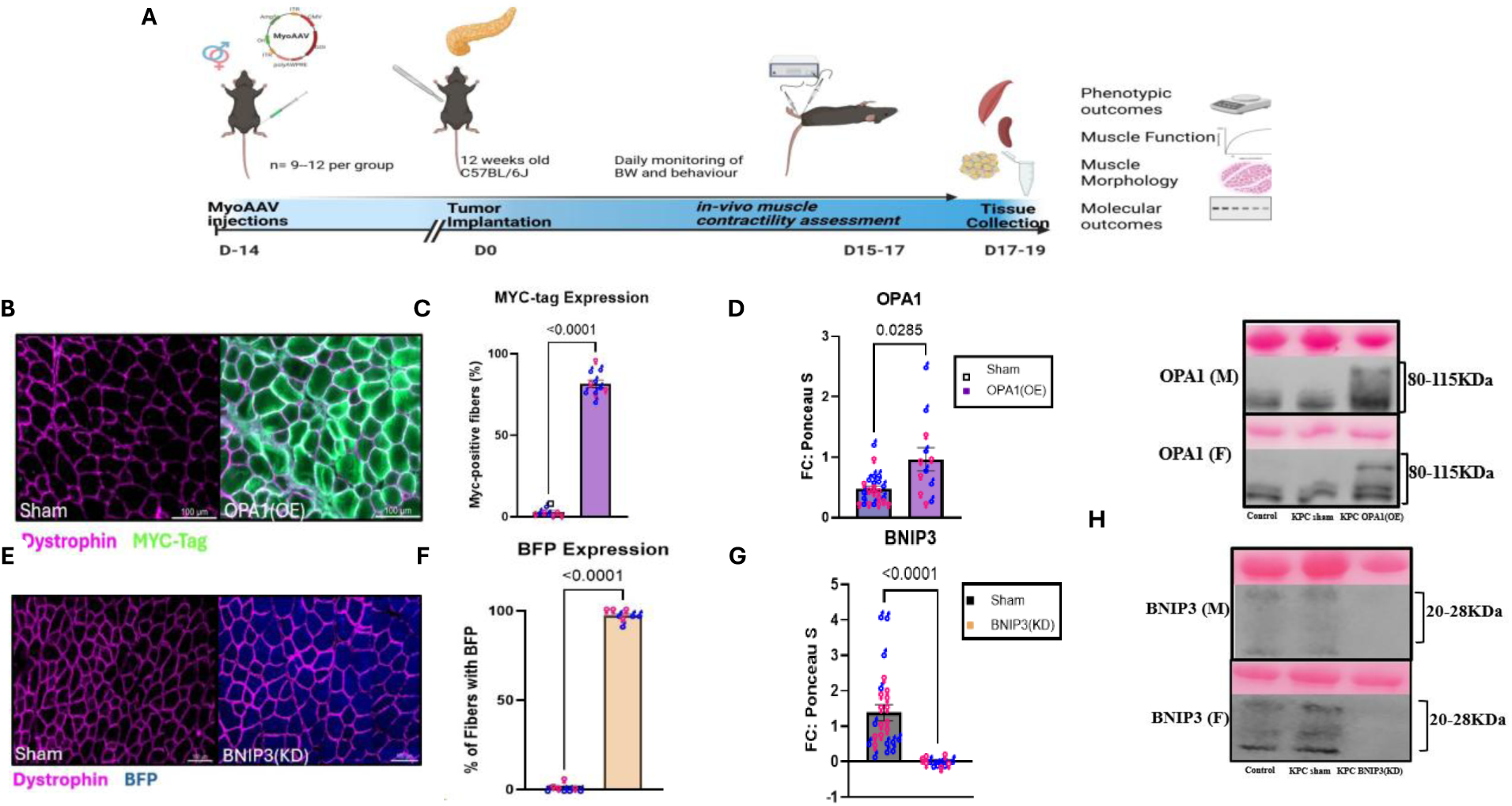
MyoAAV-mediated gene delivery effectively transduces TA muscle fibers and produces modulation of OPA1 or BNIP3 protein. (A) Experimental design and timeline for both experiments. (B) Representative immunohistochemical images of TA cross-sections stained for Myc-tag protein (green) to mark transduced fibers and dystrophin (magenta) to mark fiber borders, scale bar= 100µm. (C) the percentage of Myc-tag positive fibers per cross section in sham and OPA1(OE) muscle to indicate the transduction efficiency, (n= 8-12 per group) (D) Abundance of OPA1 protein in TA muscle expressed as fold change relative to Ponceau stain and normalized to internal control, (n= 12-24 per group). (E) Immunohistochemical representative images of muscle cross-sections showing expression of TagBFP-positive fibers in sham and BNIP3(KD) muscle. (F) Percentage of positive fibers for reporter gene (BFP) in sham and BNIP3(KD) muscle (**n)** (G) Protein quantification of BNIP3 protein in TA muscle expressed as fold change relative to Ponceau stain and normalized to internal control, (n= 12-24 per group). (H) Representative immunoblots with corresponding ponceau S stains for OPA1 (80-115 kDa) and BNIP3 (20-28 kDa) in males (M) and females (F) Lanes are C (Control), KS (KPC Sham), KO (KPC-OPA(OE)), KB (KPC BNIP3(KD)). Males and females are shown in different symbols with sham containing samples from both control and KPC Sham groups in C, D, F and G. Date shown as means ± SE with individual animals shown. Unpaired t test was performed to find the statistical difference (p≤ 0.05) and p values are shown in each panel. OE, overexpression; TA, tibialis anterior; KPC, Kras^G12D^/Trp53^R172H^/Pdx1-Cre; KD, knockdown; BFP, blue fluorescent protein.

## RESULTS

### MyoAAV effectively increases OPA1 or decreases BNIP3 expression in skeletal muscle

In order to modulate mitochondrial fusion and mitophagy in TA muscle, we selectively delivered myotropic AAV (MyoAAV) vectors to encode Myc-tagged OPA1 (OPA1 OE), or a BNIP3-targeting shRNA (BNIP3 KD) via intramuscular injection 2 weeks prior to tumor implantation. We performed whole plasmid sequencing to confirm the presence of gene of interest (GOI) for both experiments (Supplementary Figure 1).

We first confirmed transduction of the OPA1 vector by immunostaining TA cross-sections for Myc-epitope tag and dystrophin. Myc-Tag signal (green) was present in 82% of transduced fibers in OPA1 OE muscle as compared to 3% fibers signal in sham (p<0.0001; n= 8-11 per group; Fig. 1B and 1C). We also measured Opa1 protein content in targeted muscle to investigate the transduction efficacy. OPA1 protein was 2-fold higher in OPA1 (OE) group compared to non-transduced control (p= 0.02; n= 12-24 per group; Fig. 1D and 1H). We also confirmed the transduction of the shBNIP3 vector by imaging for BFP signal. BFP signal (blue) was present in ∼98% of transduced fibers in KPC BNIP3(KD) muscle as compared to Sham (Figure 1E and 1F). Additionally, representative images for all groups and sex are represented in supplemental figure 2 (S2). Conversely, BNIP3 KD resulted in ∼98% lower protein expression (p<0.0001; n=12-24 per group; Fig. 1G and H). Together, these data validate efficient myofiber transduction and yielded the target alterations to OPA1 and BNIP3 protein in skeletal muscle.

### Muscle specific OPA1 overexpression partially preserves tibialis anterior (TA) mass without altering systemic cachexia

KPC tumors produced the phenotype characteristic of this model as significant body weight loss was observed in KPC Sham and KPC OPA1(OE) group as compared to day 0 (day of tumor implantation) in both males (p= 0.03, p<0.0001 respectively; Fig. 2A) and females (p= 0.02, p= 0.01 respectively; Fig. 2G). There was no difference in tumor weights in both cancer groups (Supplementary figure S3 A and C). To validate systemic cachexia independent of our targeted tissue, we recorded the weights of plantaris muscle, spleen and gonadal fat as cachexia/wasting indices. Plantaris and gonadal fat masses were significantly lower in both cancer groups (p=0.0003, p< 0.0001, respectively) in male mice with evident splenomegaly (p<0.0001; Fig. 2B). There was no significant difference observed between KPC Sham and KPC OPA1(OE) mice relative to these tissues. The targeted TA muscle was lower in KPC Sham group (∼-15%; p=0.002) compared to control (Figure 2C). However, the KPC OPA1(OE) group was not significantly different from Control. While not significant, mean TA mass was 9% higher in KPC OPA1(OE) compared to KPC Sham, suggesting mitigation of mass loss of the targeted muscle by OPA1 overexpression.

**Figure 2:**
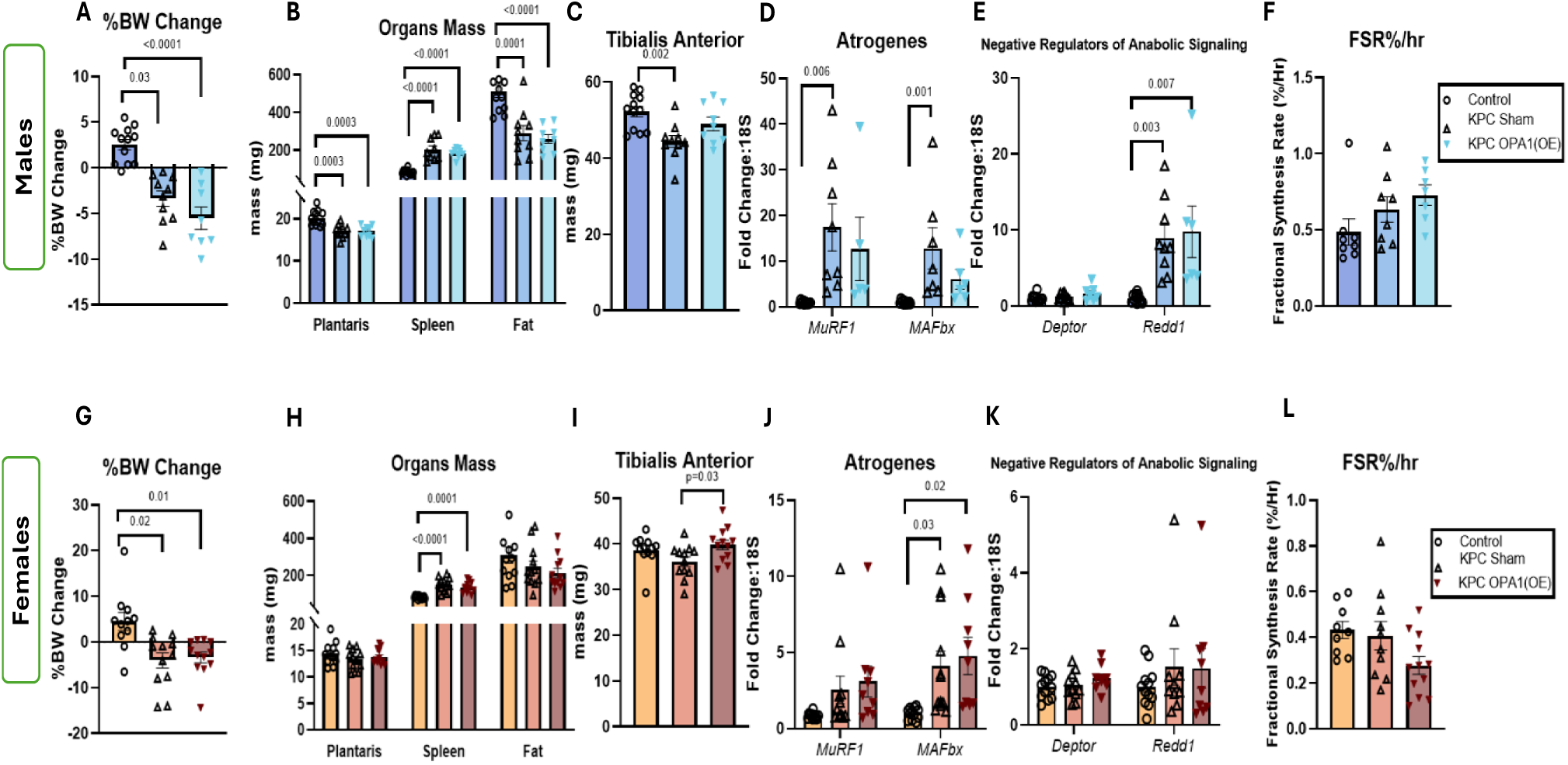
TA muscle targeted MyoAAV-mediated OPA1 overexpression partially preserves muscle mass without attenuation of systemic cachexia. (A-E) Male and (F-J) Female mice. (A and F) Percentage change in BW from initial BW (day of orthotopic KPC injections, D0) to tissue collection (D19 for males and D17 for females). (B and G) Raw wet weights of plantaris, spleen and gonadal fat at the day of tissue collection (n= 9-12per sex per group). (C and H) wet weight of TA muscle (n= 9-12per sex per group). (D and I) mRNA abundance of muscle specific E3 ubiquitin ligases *MurF1 (Trim63)* and *MAFbx (Fbxo32)* and of anabolic suppression markers *Deptor* and *Redd1 (Ddit4)* as fold change relative to *18S*. (E and J) Fractional synthesis rate (FSR) of muscle protein expressed as per hour percent of D_2_O administration over ∼ 24h prior to tissue collection (n=8-12 per group per sex). Date shown as means ± SE with individual animals shown. One-way ANOVA was performed with Tukey post hoc comparisons (p≤ 0.05) and p values are shown in each panel.

To delineate impacts on common gene programming of protein turnover, atrophy related markers were measured. Higher abundances of *MurF1* and *MAFbx* (also known as *FBXO32*) (∼17-fold change (FC), p=0.006; ∼13 FC, p= 0.001 respectively) were observed in KPC Sham compared to Controls, however, mean induction of catabolic markers observed in KPC OPA1(OE) was not significant (Fig. 2D). We next measured anabolic repressors *Deptor* and *Redd1*. There was no significant effect of cancer or treatment on *Deptor* mRNA levels, however, *Redd1* mRNA content was 8-fold greater in both cancer groups as compared to Controls (p= 0.003 (KPC Sham), p=0.007 (KPC OPA1(OE); Fig. 2E). Finally, FSR was measured as indicator of protein synthesis and was not different between experimental conditions (Fig. 2F).

In females, splenomegaly was observed in both cancer groups (KPC Sham: p<0.0001, KPC OPA1(OE): p=0.0001; Fig. 2H). While KPC did not significantly reduce TA mass (ns), TA mass of KPC OPA1(OE) group was significantly higher (∼ 9.3%) compared to KPC Sham (p= 0.03; Fig. 2I). Among catabolic markers measured, only MAFbx content was higher in response to cancer as compared to control group (Fold change of 4.1 in KPC Sham group, p= 0.03; FC of 4.6 in KPC OPA1(OE) group, p= 0.02; Fig. 2J). Fractional synthesis rate was unaffected in females as well (Fig. 2L). Raw phenotypic data is presented in supplementary table 1.

### OPA1 overexpression produces a male-specific functional protection without fatigue resistance

To further explore the effect of OPA1 overexpression, we analyzed muscle function by *in vivo* contractility assessment of dorsiflexors, with TA as the primary contributor of this muscle group. KPC Sham exhibited 33% lower torque at 10 and 20Hz as compared to healthy control (p= 0.01 and 0.002 respectively; Figure 3A). KPC OPA1(OE), however, displayed only a non-significant mean 16% lower torque compared to Control (Figure 3A). Both cancer groups displayed 33-62% lower torque than control group from 30-60Hz indicating a cancer effect unchanged by OPA1 manipulation at lower frequencies (p< 0.05). At 80 and 100 Hz, KPC Sham presented 48% (p=0.0002) and 28% (p= 0.02) lower torque compared to Control, while there was no significant difference in torque production between Control and KPC OPA1(OE). OPA1 overexpression resulted in 27% mean higher torque (p=0.07) and 23% mean higher torque (p=0.08) as compared to KPC Sham at 80 and 100 Hz stimulation frequencies respectively (Fig. 3A). As for fatigability, while a significant cancer effect was noted at 70-90 stimulations post peak torque, there was no protection observed in the cancer group with OPA1 overexpression (Fig. 3B)

**Figure 3:**
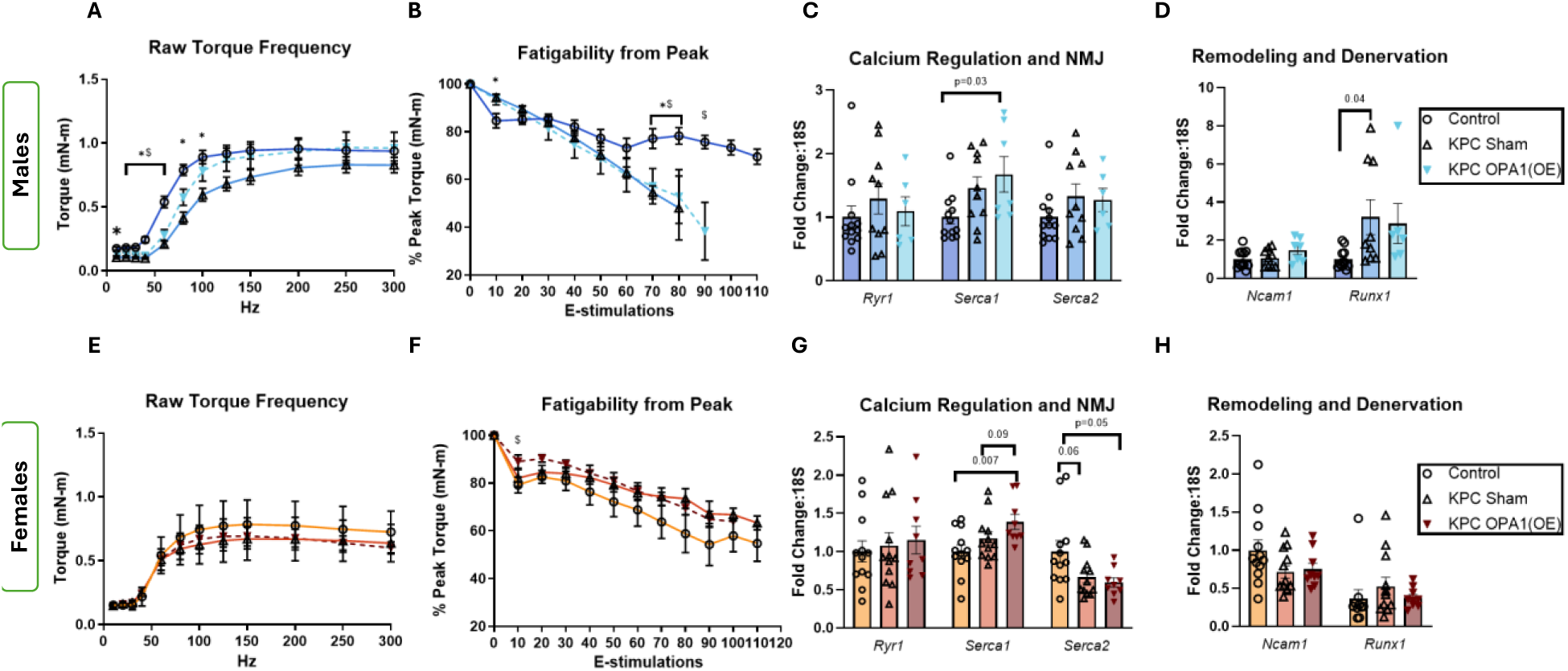
OPA1 overexpression produces a male-limited functional protection without fatigue resistance. (A-D) Male and (E-H) Female mice. (A and E) raw torque and stimulation frequency relationship of the dorsiflexion complex across 10-300 Hz range. (B and F) Fatigability expressed as % of peak torque across 120 stimulations over 120s at 40 Hz. (C and G) mRNA abundance of genes associated with calcium regulation and NMJ (*Ryr1, SERCA1* and *SERCA2*), n= 6-12 per group per sex. (G and H) mRNA abundance of remodeling genes (*Ncam1* and *Runx1)* expressed as fold change relative to 18S, n= 6-12 per group per sex. Data are described as mean ± SE. One-way ANOVA was performed with Tukey post hoc comparisons; statistical significance was identified at p ≤ 0.05 and p values are shown in each panel. *p s< 0.05, KPC Sham vs Control; #p<0.05, KPC OPA1(OE) vs KPC Sham; $p < 0.05, KPC OPA1(OE) vs Control

We next sought to assess whether improvements to torque at frequencies that produce near-max torque may be a result of improved calcium regulation and neuromuscular junction gene expression. mRNA levels of calcium regulation associated markers *Ryr1* (ryanodine receptor 1), *Serca1* (relevant to type II muscle fibers) and *Serca2* (relevant to type I muscle fibers) were measured. There was no significant effect of cancer or OPA1 overexpression observed in *Ryr1* or *Serca2* levels, however, there was a significantly higher *Serca1* gene content by fold change of ∼6 in KPC OPA1(OE) compared to Controls (p= 0.03; Fig. 3C). We then assessed gene content of remodeling and denervation markers (*Ncam1* and *Runx1*). *Runx1* levels were greater (FC:3.2; p=0.04) in KPC Sham compared to Controls, while only a more modest non-significant mean 2.8-fold elevation was observed in KPC OPA1(OE). There was no effect of cancer or OPA1 overexpression on *Ncam1* levels (Fig. 3D).

In females, there was no contractile dysfunction observed, consistent with their less overt cachectic phenotype and previous literature (11, 47, 48). Thus, there were no significant differences in torque or fatigability recorded with OPA1 overexpression (Fig. 3E-F). Despite no difference in contractile function, calcium regulatory markers were different. OPA1 overexpression exhibited significantly higher *Serca1* mRNA content compared to Control (FC:1.5, p=0.007) and a nonsignificant mean 21% greater *Serca1* mRNA content compared to KPC-Sham (p=0.09). A cancer effect was observed in *Serca2* content with lower levels in KPC Sham (FC:0.66, p=0.05) and KPC OPA1(OE) (FC:0.60; p=0.06) compared to Control (Fig. 3G). There were no significant differences in *Ryr1*, *Ncam1* and *Runx1* levels between any group thus indicating no effect of cancer or treatment.

### OPA1 overexpression leads to shift in fiber-type distribution without restoring mean fiber CSA

The TA muscle phenotype was further assessed for mean fiber cross sectional area (CSA), fiber type specific CSA, and fiber type distribution. We observed an overall lower mean CSA in male tumor-bearing groups compared to Controls (19% lower in KPC Sham, p= 0.01; 18% lower in KPC OPA1(OE), p= 0.02) with no difference between KPC Sham and KPC OPA1(OE) groups (Fig. 4B). Considering fiber-type specific CSA, we observed 30% lower CSA in type IIB fibers of KPC Sham compared to Control (p=0.002). However, OPA1 overexpression resulted in 20% greater CSA (type IIB fibers) compared to KPC Sham (Fig. 4C). Muscle specific fiber-type distribution was also assessed with significantly fewer (−13%, p= 0.001) IIB fibers in KPC Sham compared to Control, while KPC OPA1(OE) showed no difference in the IIB fiber pool compared to Control in males with 28%, more IIB fibers compared to KPC Sham (p= 0.03, KPC Sham, Fig. 4D).

**Figure 4:**
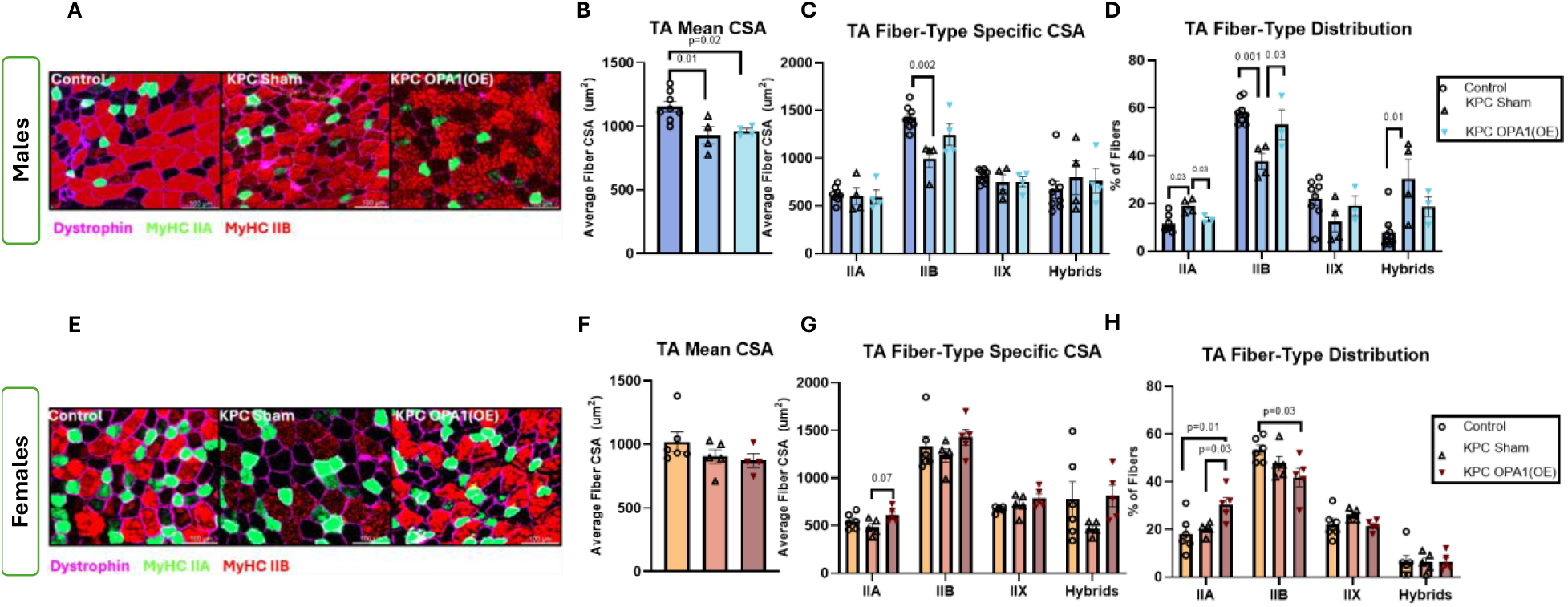
OPA1 overexpression causes the shift in fiber-type distribution toward the control phenotype without restoring mean fiber CSA. Male (A-D) and female (E-H) mice. (A and E) Representative immunofluorescence images of TA cross sections stained for dystrophin (magenta, fiber borders), MyHC IIA (green, type IIA fibers), MyHC IIB (red, type IIB fibers), unstained/negative fibers were classified as MyHC IIX and fibers positively stained for more than one isoform were classified as hybrids. Scale bar = 100um. (B and F) Mean CSA within each cross-section. (C and G) Fiber-type specific CSA for IIA, IIB, IIX and hybrid fibers. (D and H) Fiber-type specific distribution within each TA muscle cross-section expressed as the percentage of total fibers Data shown as mean ± SE. n= 4-8 per group per sex, One-way ANOVA was performed with Tukey post hoc comparisons, statistical significance was identified at p ≤ 0.05 and p values are shown in each panel. CSA, cross-sectional area; MyHC, Myosin heavy chain; TA, tibialis anterior

In females, there was no effect of cancer on TA mean CSA, consistent with the mild cachectic phenotype (Fig. 4F). However, the proportion of type IIA fibers were greater in KPC OPA1(OE) group than KPC Sham (+36%; 0.03) and Controls (+40%; p= 0.01). Type IIA fiber’s mean CSA was also 20% higher in KPC OPA1(OE) than KPC Sham although the difference did not reach statistical signific (p= 0.07) (Fig. 4H). In females, KPC OPA1(OE) had fewer type IIB fibers than Control (−23%, p= 0.03; Fig. 4H).

### OPA1 overexpression does not change oxidative phenotype of the TA muscle in either sex

We then assessed the effect of cancer or OPA1 overexpression on oxidative phenotype of the TA muscle utilizing SDH staining. The percentages of SDH positive fibers classified as light, medium or dark stained in Control were 36 ± 9%, 50 ± 6%, and 12 ± 3%, respectively. In KPC Sham group, the corresponding percentages were 34 ± 10%, 48 ± 7%, and 17 ± 3%. In KPC OPA1(OE) group these percentages were 52 ± 10%, 36 ± 9%, and 11 ± 4%, respectively (Fig. 5B). Although, there was no significant difference among groups, consistent with lack of difference in COXIV protein indicating no or very little effect of cancer on oxidative capacity of muscle in males (Fig. 5C). A similar pattern was observed in females as well. OPA1 protein levels were elevated in KPC OPA1(OE) compared to Control as well as KPC Sham animals (+55%, p= 0.05; +63%, p= 0.02 respectively). OPA1 protein levels were higher in the treated group in females by 54% (p=0.08) as compared to control group in females as well (Fig. 5G and H). There was no significant difference in DRP1 protein expression in both male and female mice (Fig. 5C and 5G respectively).

**Figure 5:**
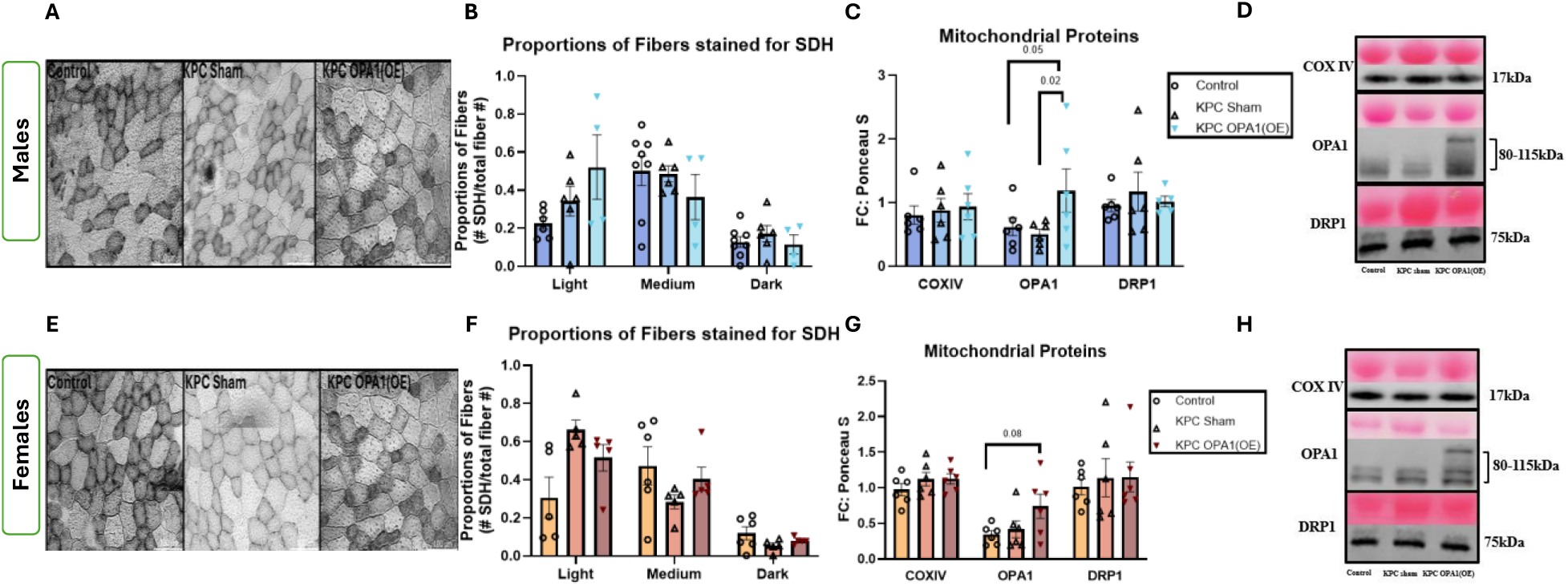
OPA1 overexpression does not change oxidative capacity of the TA muscle in either sex. Male (A-D) and female (E-H) mice. (A and E) Representative images of SDH histochemical staining of TA cross-sections, Scale bar = 100µm. (B and F) Proportions of fibers stained and classified as light, medium or dark intensity expressed as fraction of total fibers per cross-section (n= 4-8 per group per sex). (C and G) immunoblot measurements of OPA1, DRP1 and COXIV protein expressed as fold change to ponceau stain and normalized to internal control (n= 6 per group per sex). Data shown as mean ± SE. One-way ANOVA was performed with Tukey post hoc comparisons; statistical significance was identified at p ≤ 0.05 and p values are shown in each panel.

### MyoAAV mediated muscle specific BNIP3 knockdown in the TA muscle produces partial protection of muscle mass without systemic rescue and lowers basal protein synthesis in both sexes

Similar to the OPA1 cohort, tumor bearing animals displayed a cachectic phenotype with significant change in BW since day 0. KPC Sham and KPC BNIP3(KD) groups showed significant BW loss compared to Controls in males (p= 0.02, p<0.0001 respectively) and females (p= 0.002, p= 0.001 respectively) (Fig. 6A and 6G). The endpoint weights of plantaris, spleen and gonadal fat were also recorded. In males, significant cancer effect was observed in KPC Sham and KPC BNIP3(KD) groups with significant lower mass of plantaris (−15%, p=0.001; −20%, p< 0.0001, respectively), gonadal fat (−43%, p= 0.0003; −66%, p<0.0001) and evident splenomegaly (+141%, p<0.0001; +74.4%, p= 0.0006 respectively; Fig. 6B). The TA muscle was 14.9% smaller in KPC Sham animals (p= 0.07) as compared to control group; however, KPC BNIP3(KD) group was not significantly different from Control (Fig. 6C). When we looked at catabolic markers. Cancer effect was observed in MurF1 and MAFbx as MurF1 was 18 and 34-fold higher in abundance in KPC Sham (p= 0.03) and KPC BNIP3(KD) group (p<0.0001) as compared to Control (Fig. 6D). MAFbx was 15-fold higher in its gene content in KPC BNIP3(KD) as compared to Control (p= 0.0002). Similar pattern of induction with cancer was observed in anabolic suppression markers, where *Deptor* was 1.5-fold higher in abundance in KPC BNIP3(KD) group as compared to Control and *Redd1* was 8-fold higher in both groups (p= 0.002, p=0.0007 respectively; Fig. 6E). As unbalanced protein synthesis and degradation are one of the contributing factors of muscle loss in cachexia, we also sought to examine the effect of cancer and BNIP3 knockdown in targeted tissue on total protein synthesis. There was no cancer effect observed in KPC Sham group as compared to Control, but BNIP3 knockdown happened to lower fractional synthesis rate as compared to KPC Sham (−85%, p<0.0001) and Control (−81%, p= 0.002; Fig. 6F).

**Figure 6:**
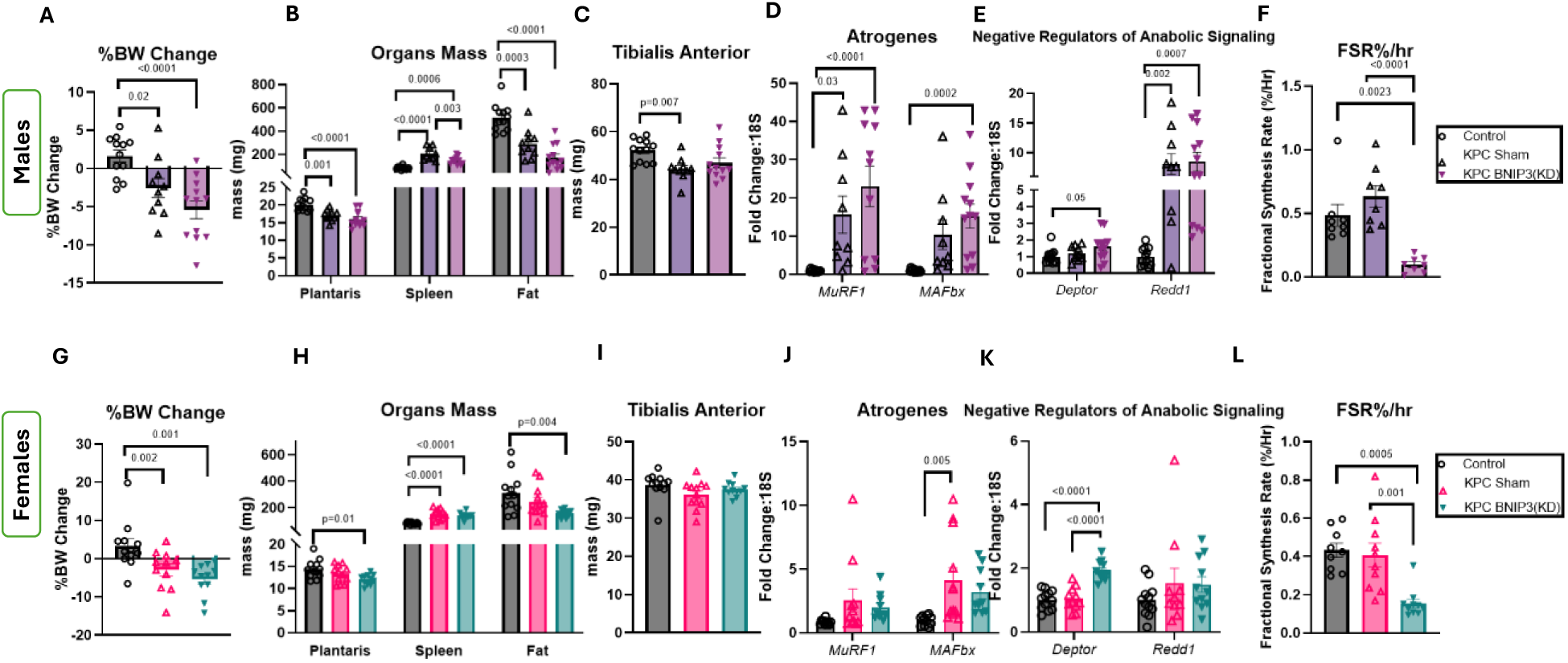
MyoAAV mediated BNIP3 knockdown in the TA muscle produces male specific partial protection of muscle mass without systemic rescue and lowers basal protein synthesis in both sexes. Male (A-E) and female (F-J) mice. (A and F) percent change in body weight between groups B and G. Raw wet weights of plantaris, spleen and gonadal fat as cachexia indicators. (C and H) Wet weight of tibialis anterior muscle. (D and I) mRNA content of *MuRF1* (*Trim63*), *MAFbx* (*Fbxo32*), *Deptor*, and *Redd1* (*Ddit4*) relative to *18S*. (E and J) TA muscle protein fractional synthesis rate expressed as percent per hour of D_2_O administration over the ∼24h preceding tissue collection Groups are Control (non-tumor bearing animals, sham vector), KPC Sham (tumor bearing animals, sham vector), KPC BNIP3(KD) (tumor bearing animals, shBNIP3 vector). Data are means ± SE with individual animals shown; n = 9–13 per sex per group. Statistical significance was identified at P ≤ 0.05 by one-way ANOVA with Tukey post hoc test; exact P values are displayed in each panel.

In females, plantaris and gonadal fat masses were lower (−16%, p=0.01; −51%, p= 0.004 respectively) in KPC BNIP3(KD) compared to Controls Figure 6H). There was significant splenomegaly in both cancer groups (+67-81%, p< 0.0001). There was no difference in TA muscle mass in response to either cancer or treatment (Fig. 6I). Among atrophy related markers, *MAFbx* mRNA content was 2-fold greater in only KPC Sham compared to Control (p= 0.005; Fig. 6J). *Deptor* mRNA content was 2-fold greater in KPC BNIP3(KD) group as compared to KPC Sham (p< 0.0001) and Control (p<0.0001; Fig. 6K). FSR was lower in females as well with the knockdown of BNIP3 compared to Control (−65%; p= 0.0005) and KPC Sham (−62%; p= 0.001; Fig. 6L).

### BNIP3 knockdown preserves torque production and fatigue resistance male tumor bearing mice

We further explored the effect of BNIP3 KD on contractile dysfunction observed in cachexia. We analyzed the *in vivo* muscle contractility with TA as the primary contributor for dorsiflexion function. KPC Sham displayed up to 62% lower torque from 10-150 Hz (p< 0.05) compared to Control. Additionally, KPC BNIP3(KD) TA muscle displayed up to 30% greater torque (p< 0.05) from 10-60 Hz. Greater torque (up to 70%; p<0.05) was observed in KPC BNIP3(KD) compared to KPC Sham from 10-100Hz frequency range (Fig. 7A). As for fatigability, there was no difference in torque between Control and KPC BNIP3(KD). KPC Sham was 11-22% more fatigued than KPC BNIP3(KD) at 60-80s of continuous contractions after peak torque. (p<0.05). There was no torque output from KPC Sham beyond 90 contractions after peak torque (Fig. 7B).

**Figure 7:**
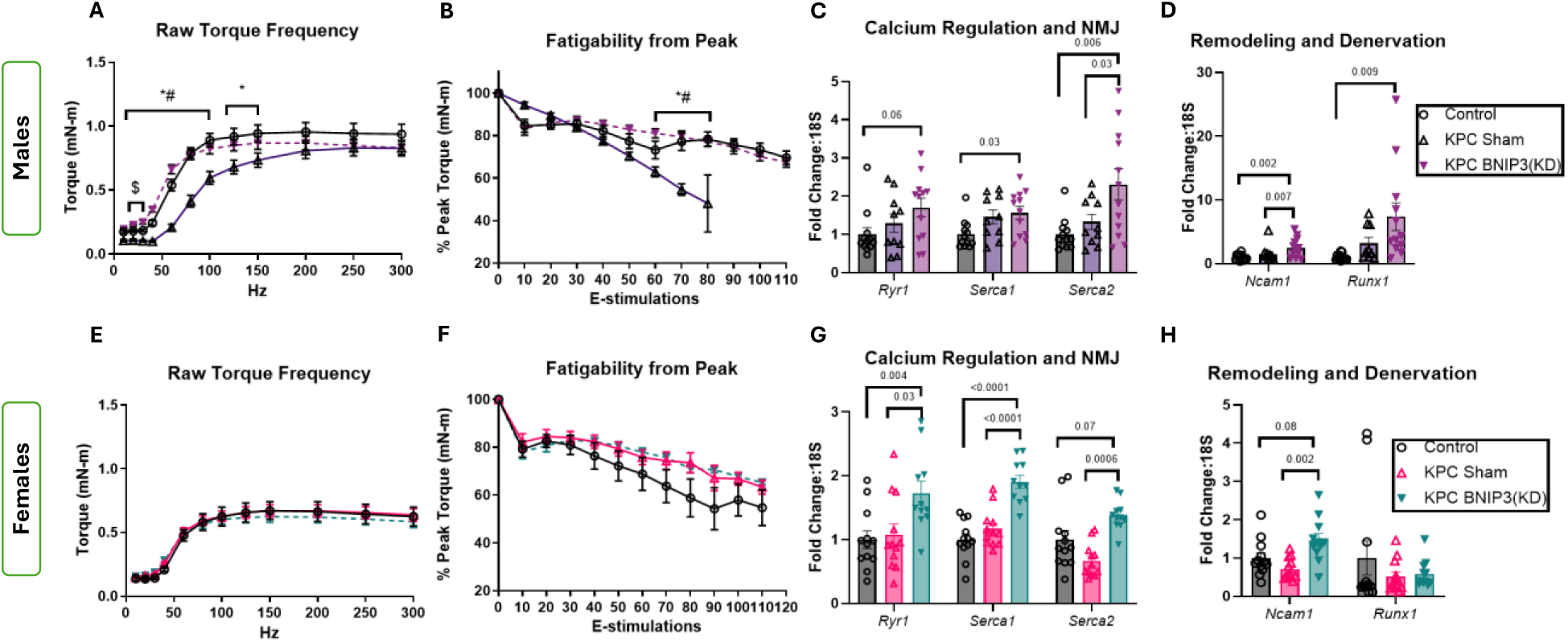
BNIP3 knockdown resulted in torque production restoration and reverses fatigability in male tumor bearing mice. Male (A-D) and female (E-H) mice. (A and E) raw torque and stimulation frequency relationship of the dorsiflexion complex across 10-300 Hz range. (B and F) Fatigability expressed as % of peak torque across 120 stimulations over 120s at 40 Hz. (C and G) mRNA abundance of genes associated with calcium regulation and NMJ (*Ryr1, SERCA1* and *SERCA2*), n= 6-12 per group per sex. (G and H) mRNA abundance of remodeling genes (*Ncam1* and *Runx1)* expressed as fold change relative to *18S*, n= 6-12 per group per sex. Data are described as mean ± SE. One-way ANOVA was performed with Tukey post hoc comparisons; statistical significance was identified at p ≤ 0.05 and p values are shown in each panel. *p < 0.05, KPC Sham vs Control; #p <0.05, KPC OPA1(OE) vs KPC Sham; $p < 0.05, KPC OPA1(OE) vs Control

We next investigated markers for calcium regulation and NMJ which contribute to force production. While Ryr1 mRNA content was not different (mean 1.6-fold change; p= 0.06), Serca1 was 1.5-fold greater compared to Control (p= 0.03). mRNA content of *Serca 2* was 1.3-fold greater in KPC BNIP3(KD) compared to KPC Sham (p= 0.0006; Figure 7C). Furthermore, mRNA content of *Ncam1* greater in KPC BNIP3(KD) compared to Control (+2.5FC; p=0.002) and KPC Sham (+1.5FC; p=0.007). Greater mRNA content of *Runx1* was only measured in KPC BNIP3(KD) compared to Control (+7FC; p= 0.009; Fig. 7D).

In females with a mild cachectic phenotype, there was no contractile dysfunction observed in cancer groups and there was no effect of BNIP3(KD) noted (Fig. 7E and 7F). However, response of calcium regulation related markers was very distinct. Greater mRNA content for *Ryr1, Serca1, Serca2* was observed in KPC BNIP3(KD) compared to Control and KPC Sham (p< 0.05; Fig. 7G). Similarly, *Ncam1* mRNA content was ∼2.6-fold greater in KPC BNIP3(KD) compared to KPC Sham (p= 0.002, Fig. 2H).

### BNIP3 knockdown partially preserves type IIB fiber cross-sectional area and proportion in male tumor-bearing mice

TA muscle was further assessed with immunofluorescence to investigate cancer and BNIP3 KD effect on muscle’s mean CSA, and fiber type specific characteristics. We observed lower mean CSA due to cancer in KPC Sham and KPC BNIP3(KD) groups (∼-20%, p= 0.01) compared to Control group (Fig. 8B). In fiber-type specific CSA, type IIB fibers CSA was lower in KPC Sham compared to Controls (−30%, p= 0.0005), while type IIB fibers CSA was more modesty 16% lower in KPC BNIP3(KD) compared to Control (p= 0.02; Fig. 8C). Considering fiber-type distribution in TA muscle, we observed 36% fewer IIB fibers in KPC Sham compared to Control (p <0.0001). KPC BNIP3(KD) showed no difference in the IIB fiber pool compared to Control, with 27% more IIB fibers compared to KPC Sham (p= 0.003; Fig. 8D).

**Figure 8:**
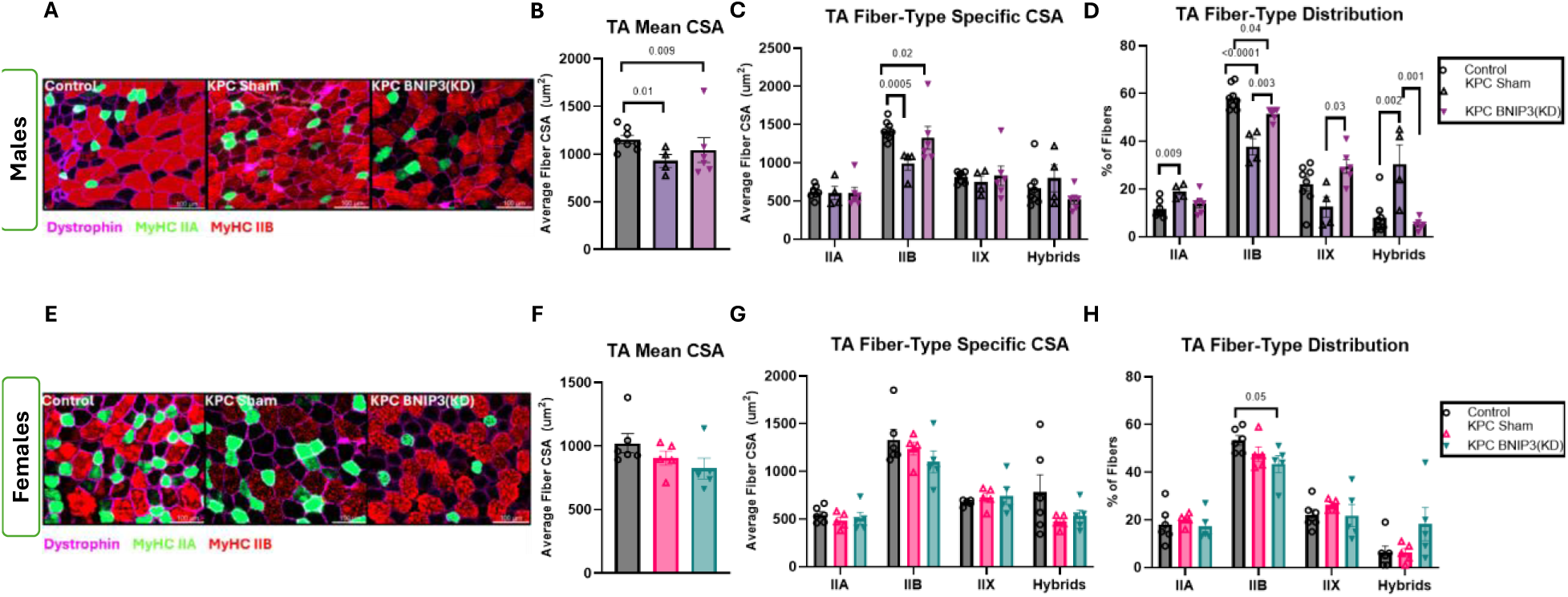
BNIP3 knockdown preserves type IIB fiber cross-sectional area and proportion in male tumor bearing mice. Male (A-D) and female (E-H) mice. (A and E) Representative immunofluorescence images of TA cross sections stained for dystrophin (magenta, fiber borders), MyHC IIA (green, type IIA fibers), MyHC IIB (red, type IIB fibers), unstained/negative fibers were classified as MyHC IIX and fibers positively stained for more than one isoform were classified as hybrids. Scale bar = 100µm. (B and F) Mean CSA within each cross-section. (C and G) Fiber-type specific CSA for IIA, IIB, IIX and hybrid fibers. (D and H) Fiber-type specific distribution within each TA muscle cross-section expressed as the percentage of total fibers Data shown as mean ± SE. n= 4-8 per group per sex, One-way ANOVA was performed with Tukey post hoc comparisons, statistical significance was identified at p ≤ 0.05 and p values are shown in each panel. CSA, cross-sectional area; MyHC, Myosin heavy chain; TA, tibialis anterior

In females, we observed no difference in mean CSA of TA muscle which coincide with mild cachectic phenotype associated with the sex (Fig. 8F and 8G). In contrast to males, KPC BNIP3(KD) females displayed 18% lower IIB proportion compared to Control (p= 0.05; Fig. 8H).

### BNIP3 knockdown does not alter oxidative phenotype but lowers DRP1 protein in males

We then assessed the effect of cancer with and without BNIP3 KD on oxidative phenotype of the targeted muscle. SDH fiber staining intensity was not different between groups in either sex (Fig. 9B and 9F) with no differences in OPA1 and COXIV protein content. While not significant, mean DRP1 protein content was 2-fold greater in KPC Sham compared to Control. DRP1 protein was 64% lower in KPC BNIP3(KD) compared to KPC Sham in males (p= 0.05; Figure 9C).

**Figure 9:**
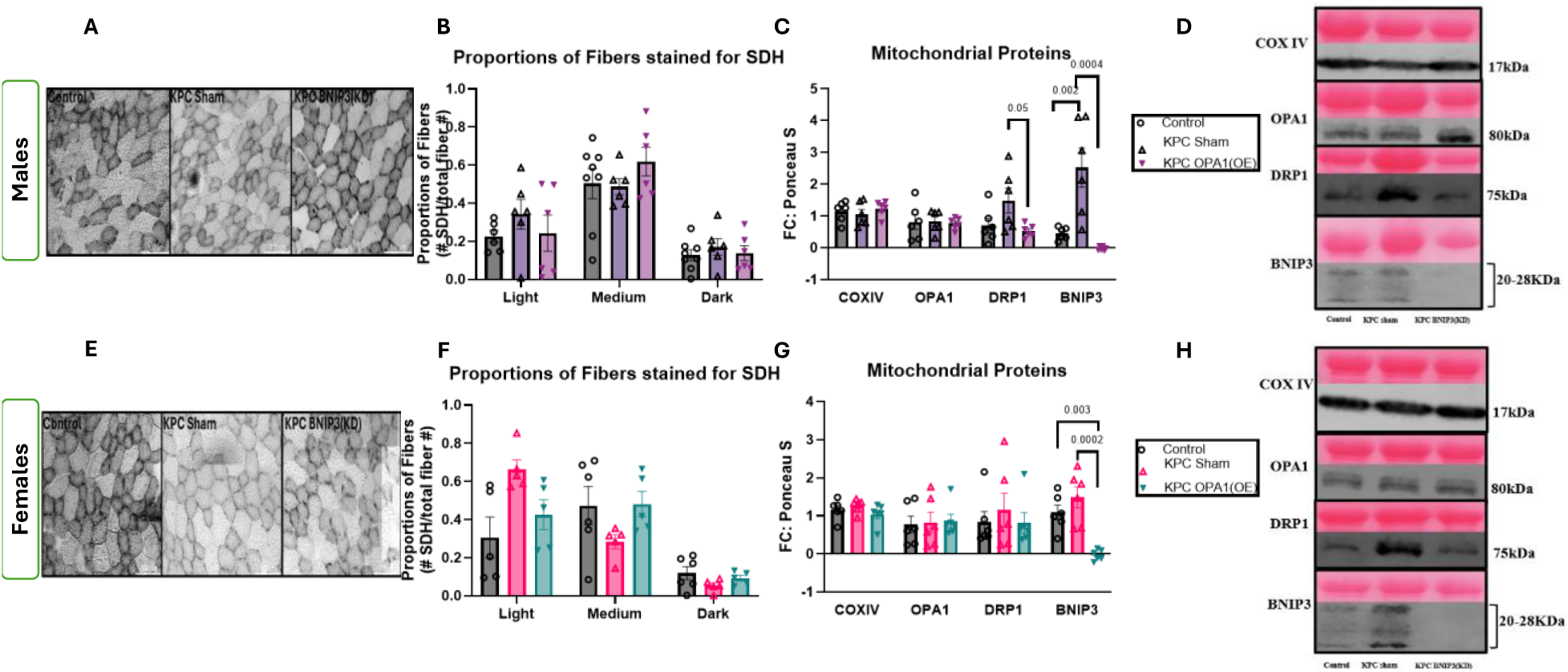
BNIP3 knockdown does not alter oxidative capacity but lowers DRP1 protein levels. Male (A-D) and female (E-H) mice. (A and E) Representative images of SDH histochemical staining of TA cross-sections, Scale bar = 100µm. (B and F) Proportions of fibers stained and classified as light, medium or dark intensity expressed as fraction of total fibers per cross-section (n= 4-8 per group per sex). (C and G) immunoblot measurements of OPA1, DRP1, BNIP3 and COXIV protein expressed as fold change to ponceau stain and normalized to internal control (n= 6 per group per sex). Data shown as mean ± SE. One-way ANOVA was performed with Tukey post hoc comparisons; statistical significance was identified at p ≤ 0.05 and p values are shown in each panel.

## DISCUSSION

Mitochondrial degeneration is believed to be an early contributing feature of atrophy with unbalanced dynamics-turnover processes in various muscle pathologies including cancer cachexia (8, 10, 49, 50). However, the efficacy of targeting mitochondrial dynamics to attenuate cachexia is not fully elucidated. Most prior studies targeting these functions have used non-tissue specific genetic models in which whole-body components are simultaneously perturbed (18, 23, 51). The present study tested whether muscle directed modifications designed to sustain mitochondrial dynamics are sufficient to improve muscle outcomes during PDAC-induced cachexia. Despite continued BW loss along with fat, splenomegaly, and wasting of non-treated muscles, MoyAAV promoting Opa1 and repressing BNIP3 each provided partial protection of targeted muscle in male mice. Most notably, both OPA1 overexpression and BNIP3 knockdown imparted significant protection of muscle contractile function in male mice. Specifically, OPA1 overexpression provided torque protection near maximal stimulation frequencies and restored the IIB fiber type proportion in males while, BNIP3 knockdown produced more pronounced protection in force production along with attenuation of fatigability and preservation in type IIB fiber type proportion. Conversely, similar to prior literature (11, 47, 52), females display a milder cachectic phenotype, representing a less responsive population to the targeted therapies applied here.

Skeletal muscle loss is most often accompanied by contractile dysfunction with reduced contractility and increased fatigability in preclinical models (8, 10, 53–55) as well as in patients (5, 6, 56–58). OPA1 overexpression improves contractile function at submaximal frequencies in male mice, in response to both muscle-directed, presented here, and whole-body/pharmacological therapies (18) employed in different pre-clinical cancer cachexia models. The data does not demonstrate direct anabolic or anti-proteolytic action as a likely primary mechanism behind these effects of OPA1. A more plausible explanation is the role of OPA1 in increasing the resilience of existing mitochondria and contractile apparatus, as it has been reported to improve respiratory efficiency and tissue resistance to injury, therefore, improving muscle quality over quantity (16, 17, 59, 60). Similar to previous reports, the contractile protection is, at least in part, supported by improved intrinsic muscle properties, likely enhanced excitation-contraction coupling rather than muscle mass gain (18, 42). Mitochondria form local networks adjacent to SR in skeletal muscle and their morphology can be disturbed through impaired calcium handling due to insufficient fusion (17) or excessive fission during repeated stimulations thus emphasizing the importance of dynamic balance (15, 61). The induction of *Serca1* after OPA1 overexpression is supportive of improved SR-reuptake of calcium, though it cannot be deduced by transcript - level changes alone. This selective response of *Serca1* induction is relevant to type IIB fibers as *Serca1* is the predominant isoform in fast-twitch skeletal muscle and supports the quick sequestration of Ca^2+^ from the cytosol to SR required for muscle contractile function (62, 63). This axis is supported by OPA1 manipulation in other cell lines and tissues. For example, OPA1 knockdown in HeLa cells, increases mitochondrial calcium uptake, while mutations in OPA1 lead to abnormal calcium homeostasis contributing to neurodegenerative diseases (64). Moreover, OPA1 heterozygous mice displayed less sensitivity of calcium accumulation to mitochondrial permeability transition pore (PTP) (65) and pharmacological targeting of mitochondria ROS through SKQ1 in C26-induced colorectal cancer and in preclinical ovarian cancer bearing mice resulted in improved contractility supported by induction of calcium handling markers (42, 66). We therefore propose that sustaining OPA1 levels preserved calcium regulation thereby improving the fast fibers activation before any measurable change in mean fiber size.

The BNIP3 intervention similarly shows therapeutic potential. Muscle restricted BNIP3 knockdown resulted in improved force production across 10-100 Hz of stimulation frequencies and attenuated fatigability to the control group, despite systemic cachexia and mean fiber atrophy. This is contrasting to global knockout of BNIP3, which exacerbated muscle atrophy and functional decline in Lewis lung carcinoma model (23), but is congruent to prior work reporting the role of muscle directed BNIP3 downregulation in attenuation of muscle loss in C26 tumor bearing mice (21). This divergence may reflect the spatial and temporal aspect of target suppression as transgenic modulation can alter development, basal mitochondrial turnover and systemic tissues. BNIP3 knockdown preserved type IIB fiber proportion along with preserved contractile function and induction of *Serca1* and *Ncam1* along with lower DRP1 levels compared to KPC Sham muscle. Lower DRP1 levels could reduce the network fragmentation which precedes muscle atrophy and enable maintenance of fast-fiber proportions (15, 61). Greater *Serca1* and *Serca2* mRNA content could improve calcium regulation and fatigue resistance (67–70), however, *Ncam1* and *Runx1* are more related to denervation and remodeling (71)*. Runx1* induction after denervation is protective against excessive autophagy (72), thus, this response may reflect an adaptive response rather than protection of the neuromuscular junction (72, 73). Our results consistent with our transgenic model are the effect of reducing mitophagy on basal protein synthesis in both sexes along with induction of *Deptor* along with atrogenes (23). This supports a model in which basal autophagy is required for myofiber integrity and removal of damaged organelles, while significant inhibition of major regulatory protein such as BNIP3 can worsen muscle loss (74). On the contrary, excessive autophagy has been reported in experimental CC and in muscle from cachectic patients (75–77). However, the degree of knockdown in our model was essentially on the same level as our global KO model (23), explaining the similar response of modulation to basal protein synthesis in the targeted muscle. Taken together, these results implicate a complicated relationship in which BNIP3 associated mitophagy induction can be harmful during cachexia, whereas prolonged suppression or near-complete deletion can compromise organelle renewal, substrate availability or anabolism.

In these experiments, female tumor bearing mice displayed a mild cachectic phenotype with very low muscle wasting and no detectable functional deficit, consistent with the previous findings that females develop a later or milder functional phenotype in several preclinical models (11, 52, 78). Sex-dependent inflammatory environment, muscle morphology or mitochondrial dynamics may change the timing of cachexia and dependence on specific quality control pathways (10, 11, 22, 23) This delayed or milder cachexia phenotype and potential sex-specific mechanistic contributions to wasting initiation creates a limitation for any rescue detection. Although OPA1 overexpression increased TA mass in tumor-bearing females and shifted the fiber type to IIA dominant muscle the overall cachexia phenotype was minimal. As type IIA fibers show relatively greater mitochondrial oxidative capacity and fatigue resistance than type IIB/IIX fibers, this change may suggest sex-dependent remodeling toward more fatigue resistance phenotype (79, 80).

Considered together, the two experiments identify balance in mitochondrial dynamics as a modifier of muscle performance independent of systemic wasting and mean fiber size. OPA1 overexpression which was previously described to reinforce the fusion-cristae architecture axis (17, 81, 82), showed modest functional protection. BNIP3 knockdown, which effects organelle turnover (83, 84), here, produced a broader male torque protection and fatigue resistance but at the cost of anabolic suppression. Although OPA1 overexpression and BNIP3 knockdown target distinct components of mitochondrial remodeling, both preserved the proportion of type IIB fibers in males and improved selected contractile outcomes without restoring mean fiber cross-sectional area. This convergence suggests maintaining fast-fiber phenotype and contractile competence may be more sensitive to mitochondrial remodeling than preventing fiber atrophy itself (42, 85, 86). Mechanistically, OPA1 overexpression may preserve cristae organization, mitochondrial network, and excitation-contraction coupling (16, 17), whereas BNIP3 knockdown may limit excessive organelle turnover and reduce DRP1-associated fragmentation (23, 84). The anabolic suppression by BNIP3 KD indicates the importance of basal mitochondrial turnover for muscle protein homeostasis. As BNIP3 is induced considerably later in temporal mapping of CC (8), a therapeutically effective approach may require partial and temporal rebalancing of mitochondrial dynamics and quality control rather than targeting one single pathway. Such preservation of muscle function could ultimately be meaningful to maintain mobility, physical independence, treatment tolerance and better quality of life in cancer patients.

## LIMITATIONS AND FUTURE CONSIDERATIONS

While this study has several strengths, including muscle-target validation of previous findings from genetic animal models, validation in the orthotopic PDAC model differing for the subcutaneously induced LLC, inclusion of both sexes as biological variable, *in-vivo* function assessment accompanied by molecular outcomes, several limitations need to be considered. We did not directly measure mitochondrial respiration, ultrastructure, reactive oxygen species, cristae architecture or damaged mitochondria (for mitochondrial turnover), relying on previous works to speculate on such features. In this study, the highly directed measure of our MyoAAV intervention led to the need to focus our assessments. Therefore, future studies should include direct measurements describing the interactive effects of mitochondrial-targeted and muscle directed approaches with wasting. OPA1 and BNIP3 intervention groups were conducted in independent and separate cohorts thus precluding the direct comparison of effect size or interaction between the two targets. Moreover, the milder cachectic phenotype observed in females may have limited detection of sex-specific effects. Future work should use inducible dosing after cachexia is established along with an additional group to investigate the combined effects of OPA1 overexpression and BNIP3 knockdown.

## CONCLUSION

Muscle targeted changes in mitochondrial dynamics improve muscle quality and contractile function in the presence of systemic cachexia in PDAC tumor bearing mice, building on our previous findings of conferred benefits against cachexia in an LLC model (18, 23). OPA1 overexpression resulted in partial, male-limited protection of TA mass and submaximal torque along with restoring IIB fiber proportion extending our transgenic findings (18) in a more translatable setting and in a different cancer type. BNIP3 knockdown conferred partial protection in muscle mass but along with broader functional benefits by restoring torque across a wide range of physiological frequencies, resisting fatigability, preserving type IIB fiber size and proportion along with lowering DRP1 levels while sustaining the same level of protein synthesis suppression as our global knockout model (23). Overall, our data show promise in targeting mitochondrial dynamics through a more clinically relevant approach in preservation of muscle function during CC. These findings also support further developments of muscle specific AAV-based approaches while also working on dose, timing and preservation of basal mitochondrial turnover as central considerations for translational relevance.

## AUTHOR CONTRIBUTIONS

**Ruqaiza Muhyudin:** Investigation, Validation, Formal analysis, Writing (original draft and Review/Editing), Visualization and Project administration **Nicole N. Noga:** Investigation and Writing (Review/Editing) **Francielly Morena:** Investigation and Writing (Review/Editing) **Sydney Hilgenbrink:** Investigation and Writing (Review/Editing) **Muskan Taori:** Investigation and Writing (Review/Editing) **Allie Ivy:** Investigation and Writing (Review/Editing) **Maryam Khaledabadi:** Investigation and Writing (Review/Editing) **Mary S. Jia:** Investigation and Writing (Review/Editing) **Christopher Nelson:** Resources, Supervision and Writing (Review/Editing) **Tyrone Washington:** Resources, Supervision and Writing (Review/Editing) **Nicholas P. Greene:** Funding acquisition, Conceptualization, Resources, Supervision, and Writing (Review/Editing)

## DATA AVAILABILITY

The supporting data will be made available upon reasonable request.

