## Supplementary Table 1 for "Mitochondrial Dynamics Modulation via MyoAAV-Mediated OPA1 Overexpression or BNIP3 Knockdown in Pancreatic Ductal Adenocarcinoma (PDAC)-Induced Cachexia"

**Supplementary Table 1: Male and Female Raw Phenotypic Table**

**Raw Weights**

| MALE | Control n=12 | KPC Sham n=10 | KPC OPA1(OE) n=9 |
|---|---|---|---|
| **Initial Body Weight (BW) (g)** | 27.43 ± 0.44 | 26.32 ± 0.57 | 28.33 ± 0.80 |
| **Final BW (g)** | 28.14 ± 0.52 | 25.45 ± 0.66* | 26.46 ± 0.71 |
| **Plantaris (mg)** | 20.17 ± 0.51 | 17.07 ± 0.49* | 17.03 ± 0.43* |
| **Tibialis Anterior (mg)** | 52.28 ± 1.36 | 44.44 ± 1.54* | 49.05 ± 1.77 |
| **Spleen (mg)** | 84.44 ± 3.74 | 204.26 ± 19.32* | 183.64 ± 7.89* |
| **Fat (mg)** | 513.03 ± 33.93 | 290.42 ± 39.59* | 259.23 ± 23.16* |
| **Tumor/Pancreas (mg)** | 181.03 ± 12.81 | 1294.84 ± 134.78* | 1549.20 ± 136.61* |

| FEMALE | Control n=12 | KPC Sham n=12 | KPC OPA1(OE) n=12 |
|---|---|---|---|
| **Initial Body Weight (BW) (g)** | 20.59 ± 0.41 | 21.89 ± 0.45 | 21.63 ± 0.39 |
| **Final BW (g)** | 21.39 ± 0.44 | 20.98 ± 0.46 | 20.88 ± 0.29 |
| **Plantaris (mg)** | 14.37 ± 0.60 | 13.28 ± 0.56 | 13.77 ± 0.42 |
| **Tibialis Anterior (mg)** | 38.70 ± 0.96 | 36.13 ± 1.05 | 39.86 ± 1.05# |
| **Spleen (mg)** | 82.88 ± 2.42 | 151.57 ± 10.59* | 134.97 ± 8.32* |
| **Fat (mg)** | 309.33 ± 42.68 | 245.87 ± 32.54 | 212.22 ± 25.25 |
| **Pancreas/ Tumor (mg)** | 138.66 ± 7.54 | 891.88 ± 76.16* | 819.13 ± 63.54* |

Values are mean ± SEM. *p<0.05 vs Control; #p<0.05 vs KPC Sham; $p<0.05 vs KPC OPA1(OE) One-Way ANOVA (Tukey-Kramer)

**Supplementary Table 2: Male and Female Raw Phenotypic Table**

**Raw Weights**

| MALE | Control n=12 | KPC Sham n=10 | KPC BNIP3(KD) n=12 |
|---|---|---|---|
| **Initial Body Weight (BW) (g)** | 27.43 ± 0.44 | 26.32 ± 0.57 | 26.72 ± 0.37 |
| **Final BW (g)** | 28.14 ± 0.52 | 25.45 ± 0.66* | 25.31 ± 0.57* |
| **Plantaris (mg)** | 20.17 ± 0.51 | 17.07 ± 0.49* | 16.07 ± 0.65* |
| **Tibialis Anterior (mg)** | 52.28 ± 1.36 | 44.44 ± 1.54* | 47.06 ± 2.02 |
| **Spleen (mg)** | 84.44 ± 3.74 | 204.26 ± 19.32* | 147.29 ± 9.01*# |
| **Fat (mg)** | 513.03 ± 33.93 | 290.42 ± 39.59* | 171.61 ± 30.86* |
| **Tumor/Pancreas (mg)** | 181.03 ± 12.81 | 1294.84 ± 134.78* | 1316.39 ± 63.70* |

| FEMALE | Control n=12 | KPC Sham n=12 | KPC BNIP3(KD) n=11 |
|---|---|---|---|
| **Initial Body Weight (BW) (g)** | 20.59 ± 0.41 | 21.89 ± 0.45 | 21.61 ± 0.30 |
| **Final BW (g)** | 21.39 ± 0.44 | 20.98 ± 0.46 | 20.43 ± 0.23 |
| **Plantaris (mg)** | 14.37 ± 0.60 | 13.28 ± 0.56 | 12.02 ± 0.36* |
| **Tibialis Anterior (mg)** | 38.70 ± 0.96 | 36.13 ± 1.05 | 37.50 ± 0.56 |
| **Spleen (mg)** | 82.88 ± 2.42 | 151.57 ± 10.59* | 139.67 ± 6.96* |
| **Fat (mg)** | 309.33 ± 42.68 | 245.87 ± 32.54 | 156.90 ± 8.53* |
| **Tumor/Pancreas (mg)** | 138.66 ± 7.54 | 891.88 ± 76.16* | 798.25 ± 83.61* |

Values are mean ± SEM. *p<0.05 vs Control; #p<0.05 vs KPC Sham; $p<0.05 vs KPC BNIP3(KD) One-Way ANOVA (Tukey-Kramer)

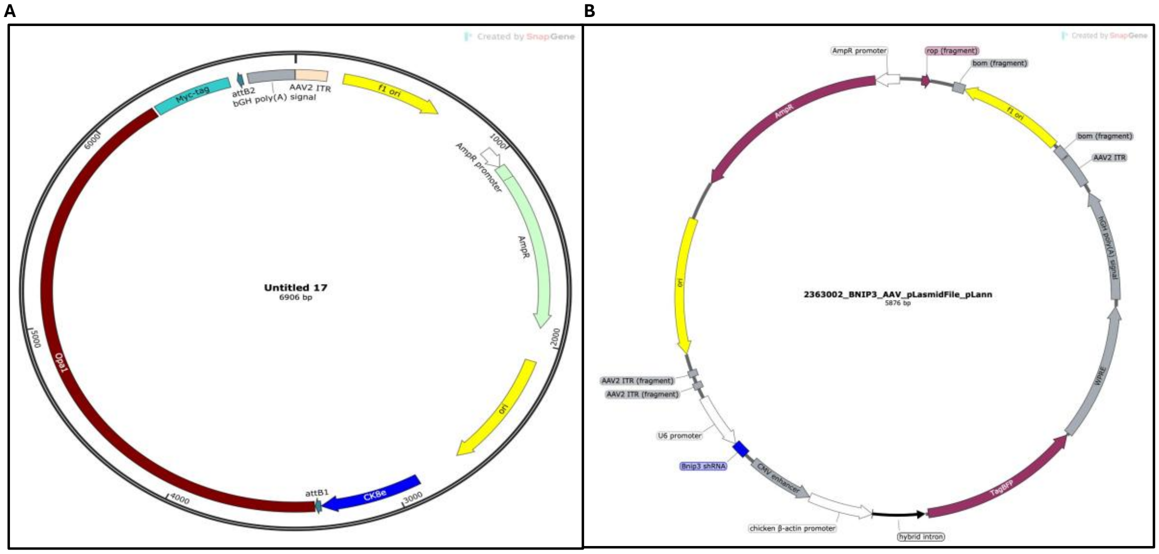

**Supplementary Figure S1: AAV plasmid constructs for OPA1 overexpression and BNIP3 knockdown** (A) Plasmid map of the AAV construct used to overexpress OPA1 in skeletal muscle. (B) Plasmid map of the AAV construct for BNIP3 knockdown encoding shBNIP3

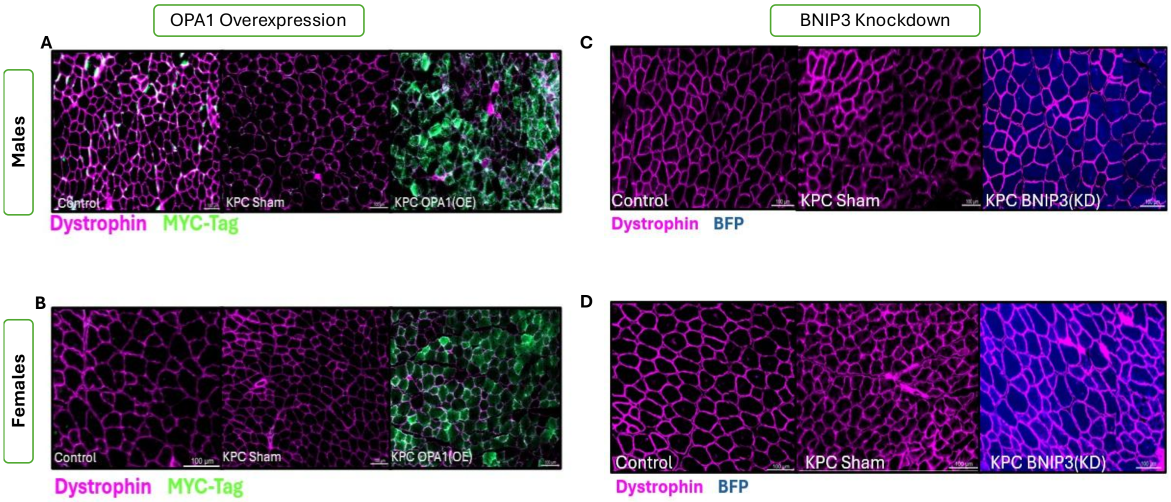


**Supplementary Figure S2: Immunofluorescence (IF) validation of AAV-mediated reporter gene transduction in muscle fibers.** Representative immunofluorescence images of tibialis anterior cross-sections from (A) male and (B) female Control, KPC Sham, and KPC OPA1(OE) mice. Dystrophin staining (magenta) for myofiber borders, while MYC-tag staining (green) identifies myofibers expressing the 13×Myc-tagged OPA1 construct. Scale bar, 100 µm. (C) male and (D) Control, KPC Sham, and KPC BNIP3(KD) mice. Dystrophin staining (magenta) for myofiber borders and blue for BFP reporter expression.

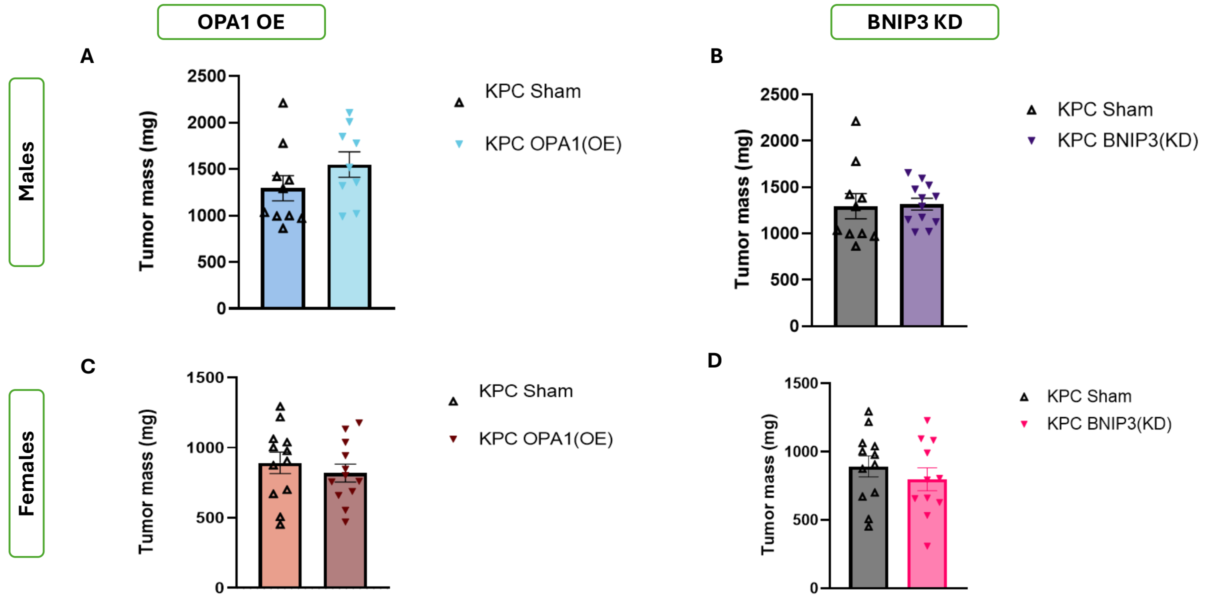


**Supplementary Figure S3: OPA1 overexpression and BNIP3 knockdown do not alter tumor mass.** Tumor mass was assessed in male and female tumor-bearing mice from the OPA1 overexpression and BNIP3 knockdown experiments. (A) Tumor mass in male KPC Sham (n=10) and KPC OPA1(OE) (n=9) mice. (B) Tumor mass in male KPC Sham (n=10) and KPC BNIP3(KD) (n=12) mice. (C) Tumor mass in female KPC Sham (n=12) and KPC OPA1(OE) (n=12) mice. (D) Tumor mass in female KPC Sham (n=12) and KPC BNIP3(KD) (n=11) mice. Bars represent mean ± SEM, and individual symbols represent individual animals. Groups were analyzed separately within each experiment and sex using one-way ANOVA. No significant differences were detected between KPC Sham and the respective intervention group (P>0.05).
